# A Functional Placenta-On-Chip Model For Maternal–Fetal Transport

**DOI:** 10.64898/2026.02.22.706965

**Authors:** Anshul Bhide, Sourav Mukherjee, Kinjalka Ghosh, Abhijit Majumder, Deepak Modi

## Abstract

The human placenta functions as a highly specialised barrier that integrates trophoblast differentiation, endocrine activity, and regulated transport of molecules to sustain fetal development. Experimental interrogation of placental barrier function remains challenging due to limited access to human placental tissue and the complexity of existing in vitro models. Here, we report a static, two-chamber placenta-on-chip platform designed to recapitulate key structural and functional attributes of the human placental barrier within an experimentally accessible format. The device design prioritises open maternal compartmentalisation and diffusion-dominated transport, reflecting the haemochorial nature of human placentation. It also remains compatible with standard multi-well plate formats for parallel experimentation. In this two-chambered device, separated by an extracellular matrix-loaded/coated microporous membrane, the trophoblast supports trophoblast syncytialisation, sustained β-human chorionic gonadotropin secretion, and selective barrier function. The engineered barrier restricts macromolecular transport while permitting controlled diffusion of small solutes. Glucose transport across the device is strongly dependent on cellular configuration, with inclusion of the endothelial layer significantly modulating nutrient flux and yielding fetal-to-maternal glucose ratios comparable to those reported in vivo. The platform further supports directional transfer of urea from the fetal to the maternal compartment, demonstrating bidirectional metabolite exchange relevant to placental waste clearance. Under hyperglycemic conditions, glucose transport across the barrier increases without evidence of barrier breakdown, indicating sensitivity to metabolic perturbation. This scalable design of a placenta-on-chip platform provides a robust framework for studying placental transport, metabolic regulation, and barrier integrity. It offers broad application in placental biology, pregnancy-associated pathologies, and screening for pregnancy-safe drugs.

## 1. Introduction

The human placenta is a transient yet indispensable organ essential for appropriate fetal development. It is a mediator of the exchange of nutrients, gases, and metabolic waste between the maternal and fetal circulations, while simultaneously acting as a selective biochemical and immunological barrier [1–3] . Structurally, this exchange is orchestrated by a highly specialised interface comprising a multinucleated syncytiotrophoblast layer in direct contact with maternal blood and an underlying fetal endothelium [2,3]. Together, these barriers regulate transport processes that are tightly tuned to gestational demands. Perturbations in placental barrier function or nutrient transport are central to several pregnancy-associated disorders, including intrauterine growth restriction, pre-eclampsia, and gestational diabetes mellitus (GDM) [4,5]. Despite its critical role in pregnancy outcome, direct experimental interrogation of human placental function is challenging due to ethical constraints, limited access to gestational tissue, and the dynamic nature of placental differentiation [6–8].

In vitro models have long been used to study placental transport and endocrine function. Conventional approaches, including trophoblast monolayers, transwell-based co-cultures, and placental explants, have provided important insights but remain limited in their ability to capture the spatial organisation and functional integration of the maternal–fetal interface [8–10]. Animal models, although valuable, have substantial interspecies differences in placentation, trophoblast differentiation, and transporter expression, which restrict their translational relevance to human pregnancy [11]. These challenges have driven the development of micro engineered placenta-on-chip platforms that combine human cell types with defined physical architectures to better approximate in vivo placental structure and function [10,12]

Several placenta-on-chip models have been reported for modelling healthy and diseased pregnancies, enabling drug testing, studying nutrient/toxin transfer, and understanding pregnancy complications. However, these employ perfusion-based microfluidic designs to study drug transfer and transport kinetics under dynamic flow conditions (Table 1) [13–33]. While these systems represent an important technological advance, they often rely on complex fabrication strategies, specialized equipment, and continuous flow operation, which can limit scalability and broader adoption [32]. In most platforms, maternal perfusion is implemented through enclosed microchannels that do not fully reflect the diffusion-dominated open-circulation environment of the intervillous space. Moreover, a significant proportion of existing platforms priorities either permeability or pharmacokinetic measurements, which comparatively puts less emphasis on trophoblast functional maturation, syncytialisation, and endocrine activity that are the hallmark features of the human placental barrier that are central to maternal–fetal communication (Table 1). Therefore, there is a need for experimentally accessible placental models that capture key features of the open maternal circulation, while emphasizing functional trophoblast differentiation and barrier behaviour under diffusion-dominated conditions, and yet remaining compatible with standard cell culture workflows.

Here, we sought to develop a scalable and experimentally accessible placenta-on-chip platform that captures key structural and functional features of the human maternal–fetal interface under diffusion-dominated conditions. Specifically, the objective was to establish a bilayer trophoblast–endothelial barrier capable of supporting trophoblast differentiation, endocrine activity, and selective transport of solutes within a static architecture compatible with standard cell culture workflows. By prioritising open maternal compartmentalisation and functional barrier behaviour over microfluidic complexity, this platform is intended to enable reproducible, parallel assessment of placental transport and barrier function.

## 2. Materials and Methods

### 2.1 Device design and fabrication

The placenta-on-chip device was designed as a two-chambered platform to spatially segregate maternal and fetal compartments separated by a semipermeable membrane. Device components were designed using AutoCAD and fabricated from acrylic sheets using a CO₂ laser cutting and engraving system (SIL Laser Cutting and Engraving Machine; AccuCut 1212/6090).

Circular reservoirs of desired dimensions were cut from thick (6 mm) acrylic sheets at 85 % laser power and a speed of 8 mm s-1. Thin acrylic sheets (1 mm) were laser-cut to generate circular lids containing two ports, which served as inlet and outlet access points. All acrylic components were cleaned by immersion in 70 % ethanol for 2 h, rinsed thoroughly with sterile Milli-Q water, and air-dried at room temperature.

The two reservoirs were bonded across a track-etched polycarbonate membrane (Whatman Cyclopore, 1 µm pore size, 10 mm diameter) using a PDMS stamp-bonding approach. Briefly, polydimethylsiloxane (PDMS; Sylgard 184, Dow Corning) was prepared by mixing the base and curing agent at a 10:1 (w/w) ratio, degassed under vacuum, and spin-coated onto a glass slide (50 × 75 mm) at 2000 rpm for 20 s. The acrylic components were gently pressed onto the PDMS-coated surface to transfer a thin adhesive layer and assembled with the membrane positioned centrally between the two chambers. The assembled device was cured at 60 °C overnight to achieve permanent bonding. Subsequently, the inlet/outlet lids were bonded using the same PDMS stamping procedure and cured under identical conditions. The final devices were compatible with standard six-well plates, enabling parallel experimentation (Figure 2).

### 2.2 Cell Culture

BeWo human choriocarcinoma trophoblast cells and HTR8-SVneo cells (both from ATCC), were cultured in DMEM/F12 (1:1) medium supplemented with 10 % fetal bovine serum (FBS) and 1 % antibiotic–antimycotic solution [34]. Human umbilical vein endothelial cells (HUVECs) were obtaiend commercially (HiMedia) maintained in HiEndoXL endothelial cell expansion medium (HiMedia) supplemented with 1% antibiotic–antimycotic solution. All cells were cultured at 37 °C in a humidified incubator with 5 % CO₂.

Cells were passaged at 75–80 % confluence using TrypLE™ Express. Cell suspensions were centrifuged at 1200 rpm for 5 min. Cells were resuspended in fresh medium and counted using a hemocytometer.

### 2.3 Co-culture in the device

Before cell seeding, the assembled devices were sterilised under ultraviolet light for 60 min (30 min per side). The membrane was coated with 2 % (v/v) Matrigel diluted in basal DMEM/F12 and incubated at 37 °C for 2 h. Excess Matrigel was removed, and chambers were washed with DPBS.

BeWo cells were seeded into the upper (maternal) chamber at a density of 1.5 × 10⁵ cells per device and allowed to attach overnight. Trophoblast syncytialisation was induced by treating BeWo cultures with forskolin (50 µM) for 48 h [35]. Following trophoblast differentiation, devices were inverted, and HUVECs were seeded into the lower (fetal) chamber at a density of 1 × 10⁵ cells per device. Devices were incubated overnight to allow endothelial attachment before downstream assays.

### 2.4 Cell viability and immunofluorescence staining

For viability assessment, devices were disassembled, and membrane were stained with Calcein-AM (1:2000) to label live cells and Hoechst 33342 (1:5000) to label nuclei. Imaging was done using an EVOS FL-Auto fluorescence microscope.

For spatial localisation and layer verification, BeWo and HUVECs were pre-labelled with CellTracker™ Red CMTPX (10 µM) and CellTracker™ Green CMFDA (10 µM), respectively, prior to seeding. Following attachment, membrane was removed and imaged using an Operetta CLS high-content screening system.

For immunostaining, cells were fixed with 4 % paraformaldehyde for 15 min and blocked with 1 % bovine serum albumin. Primary antibodies against E-cadherin (1:50) and VE-cadherin (1:200) were incubated overnight at 4 °C. Alexa Fluor 568 and Alexa Fluor 488 secondary antibodies (1:1000) were used for visualization, and nuclei were counterstained with DAPI [33,34]. Imaging was performed using Operetta CLS system. Three-dimensional reconstructions were generated using Harmony 5.0 software.

### 2.5 Assessment of on-chip trophoblast differentiation and hormone secretion

Syncytialisation was evaluated by immunofluorescence analysis of E-cadherin localization and multinucleated cell morphology. Initial β-human chorionic gonadotropin (β-hCG) secretion was assessed qualitatively using a commercial pregnancy detection kit following the manufacturer’s instructions.

Quantitative measurement of β-hCG levels was performed using a chemiluminescent microparticle immunoassay (CMIA; Abbott). Spent medium samples were processed according to the manufacturer’s protocol, and luminescence signals were measured using a clinical immunoassay analyser. β-hCG concentrations were calculated from instrument-generated calibration curves.

### 2.6 Barrier permeability assays

Barrier integrity was evaluated using fluorescein isothiocyanate (FITC; 376 Da) and FITC–dextran (70 kDa). Tracers were added to the maternal chamber at a concentration of 0.1 mg/mL[19,27]. Samples (1 µL) were collected from the fetal chamber at defined time points (0, 0.5, 1, 2, 4, and 8h). Fluorescence intensity was measured using a Qubit 4 fluorometer (excitation 485 nm, emission 520 nm). Quantification was done using calibration curves generated from serial dilutions of FITC and FITC–dextran standards, ensuring that all measurements were within the linear detection range (Supplementary Figure 3). Fluorescence values were normalised to baseline (T_0_) before analysis.

### 2.7 Glucose and urea transport assays

For glucose transport studies, maternal chambers contained DMEM/F12 with ∼17.5 mM glucose, while fetal chambers contained ∼5.5 mM glucose. Samples (2 µL) were collected from the fetal chamber at defined intervals and analysed using a handheld glucose meter (OneTouch Select Plus). The use of a glucometer enabled measurements from microlitre-scale samples, allowing repeated sampling from the same device for kinetic analysis while minimising perturbation and inter-device variability. To validate the measurements, a subset of samples was also analysed using a standard enzymatic hexokinase-based assay. Transport kinetics were assessed under acellular, BeWo monoculture, BeWo–HUVEC co-culture, and BeWo–HTR8-SVneo co-culture conditions. Hyperglycemic conditions were simulated by increasing glucose concentrations in the maternal chamber to ∼35 mM. Parallel acellular conditions were included as controls.

For urea transport assays, urea (11 mM) was added to the fetal chamber, and samples were collected from both chambers at 0 and 8h. Parallel acellular conditions were used as controls. Urea concentrations were quantified using a commercially available enzymatic assay kit (Beckman Coulter OSR6134) following the manufacturer’s instructions.

### 2.8 Apparent permeability coefficient (P_app)

The apparent permeability coefficient (P_app) was calculated from the accumulation of solute in the receiver chamber over time. Receiver concentrations were measured at defined time points, and the transport rate (dQ/dt) was determined using dQ/dt=(ΔC×V)/Δt, where V is the receiver volume, and ΔC is the concentration difference between 2 time points in the fetal chamber. P_app was then calculated as P_app=(dQ/dt)/(A×C_0_), where A is the effective membrane area, and C_0_ is the concentration difference between the initial maternal media concentration and fetal media concentration. All P_app values are reported in cm/s and were calculated per device replicate.

### 2.9 Glucose transport dynamics using integrated and peak-based parameters

Glucose transport profiles were analyzed using non-compartmental analysis (NCA) based on time-point measurements taken at t = 0, 0.5, 1, 2, 4, and 8h.

**Area Under the curve (AUC) (Total Transport)**: To quantify the total extent of glucose transport over the 8h experimental period (AUC_0-8h_), the linear trapezoidal rule was employed. The area between consecutive time points was calculated as the average of the two rate measurements multiplied by the time interval, according to

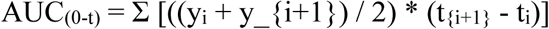

Where y_i_ represents the concentration at time t_i_. This method assumes a linear change in transport rate between sampling points. [36]

**Peak Kinetics (C_max_, R_max_, and T_max_):** Peak transport metrics were determined through direct observation of the experimental dataset, following EMA bioequivalence guidelines.[37]

*Maximum Concentration (C_max_)* was determined in accordance with ICH M13A guidelines, the highest single concentration value recorded across the sampling points was reported as C_max._

*Peak Rate (R_max_):* The rate of transport for each sampling interval was calculated as the change in concentration divided by the time interval (ΔC/Δt).

*Time to Peak (T_max_):* The specific sampling time t at which C_max_ was observed.

**Glycaemic Variability and Stability (MAGE, TIR):** To assess the stability of the transport profile, two variability metrics were calculated.

*Mean Amplitude of Glycaemic Excursions (MAGE):* MAGE was calculated to quantify the magnitude of significant transport fluctuations, excluding minor signal noise. Following the method of Service et al., excursions were defined as the absolute difference between a peak and a nadir. An excursion was included in the calculation only if its amplitude exceeded one standard deviation (>1 SD) of the device’s mean rate.[38]

MAGE = (1/k) * Σ (Peak_j_ - Nadir_j_) (calculated only for excursions > 1 SD)

*Time in Range (TIR):* TIR was defined as the percentage of the total experimental duration (8h) during which glucose levels remained within the consensus therapeutic window of 70–180 mg/dL. Since measurements were discrete, linear interpolation was used to determine the precise time coordinates where the transport curve intersected the upper (180 mg/dL) and lower (70 mg/dL) thresholds. [39]

### 2.9 RNA extraction, cDNA synthesis, and real-time PCR

RNA extraction was done using RNeasy mini kit (Qiagen, Germany). cDNA was synthesized using the High-Capacity cDNA synthesis kit (Applied Biosystems, ThermoFisher Scientific, MA, USA). Real-time PCR was performed using Eva Green chemistry (BioRad Laboratories Inc., CA, USA) in the CFX96 Real-Time PCR System (BioRad) as detailed previously [40]. PCR was carried out in duplicates for all biological replicates. Gene expression was normalized to 18S and data were analyzed using the Pfaffl method. The primer sequences and their annealing temperatures are given in Supplementary Table 2.

### 2.10 Statistical analysis

All experiments were performed with at least three independent biological replicates. Data are presented as mean ± standard deviation. Statistical analyses were conducted using two-tailed Student’s t-tests or one-way analysis of variance (ANOVA) with Tukey’s multiple group comparison procedure as appropriate. Statistical significance was defined as p < 0.05.

## 3. Results

### 3.1 Design and fabrication of a two-chamber placenta-on-chip device

To recreate the spatial organisation of the human placental barrier, we developed a static, two-chamber placenta-on-chip device comprising distinct maternal and fetal compartments separated by a semipermeable membrane (Figure 1A). The design was guided by the need to model an open maternal compartment analogous to the intervillous space, while maintaining a physically defined interface with the fetal endothelial layer (Figure 1B).

**Figure 1.**
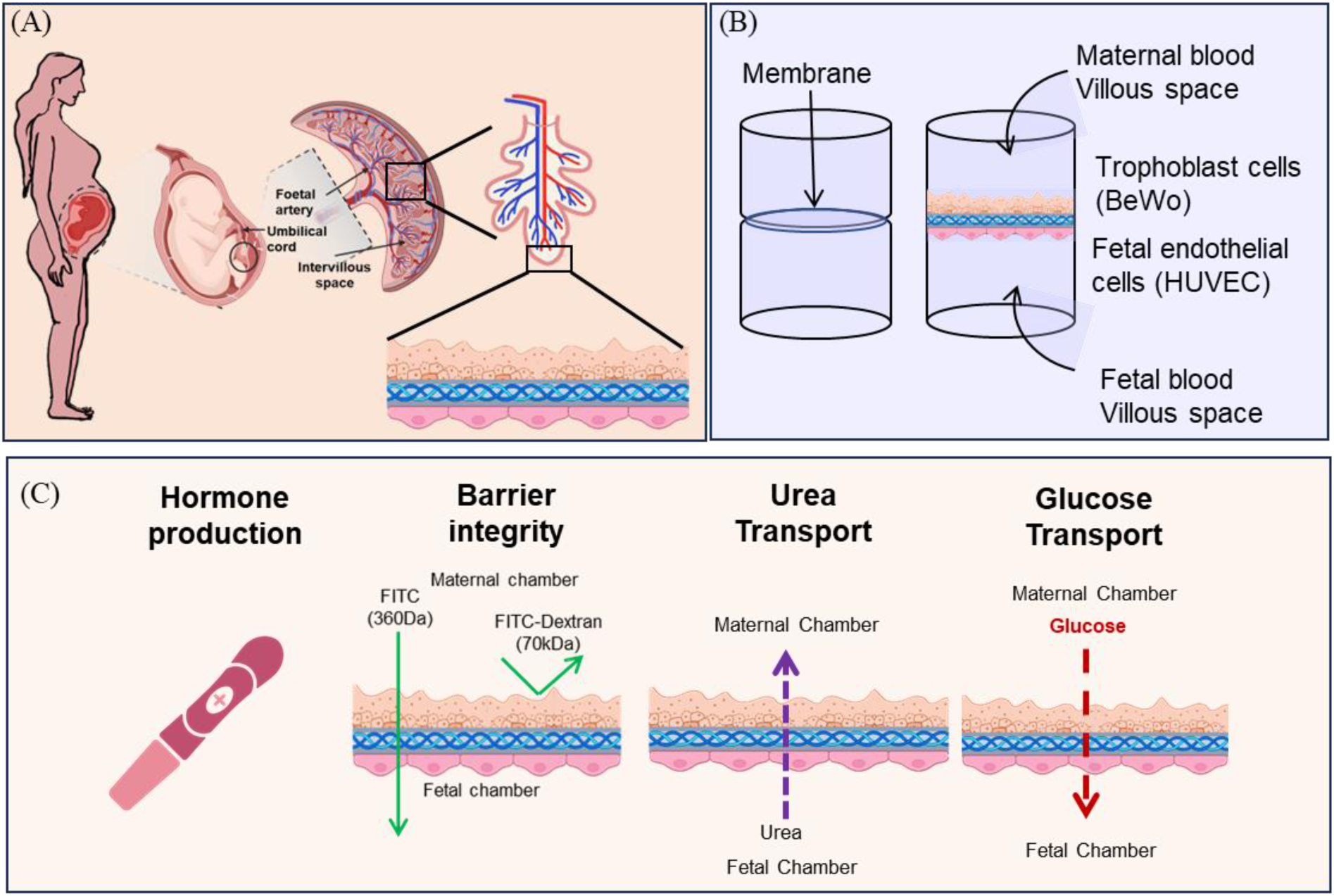
Device schematic. (A) Schematic of placental anatomy and fetal-matemal interface showing a cross-section of a pregnant woman, fetus, and placenta, highlighting the intervillous space, umbilical cord, fetal artery and the site of barrier (B) Diagram of co-culture system depicting the assembly: upper compartment (maternal villous space) seeded with trophoblast cells (BeWo cells). lower compartment (fetal villous space) seeded with Endothelial cells (HUVECS), separated by a porous membrane to mimic trophoblast and endothelial layers. (C) Overview of functional assays performed using the placenta-on-chip platform

**Figure 2.**
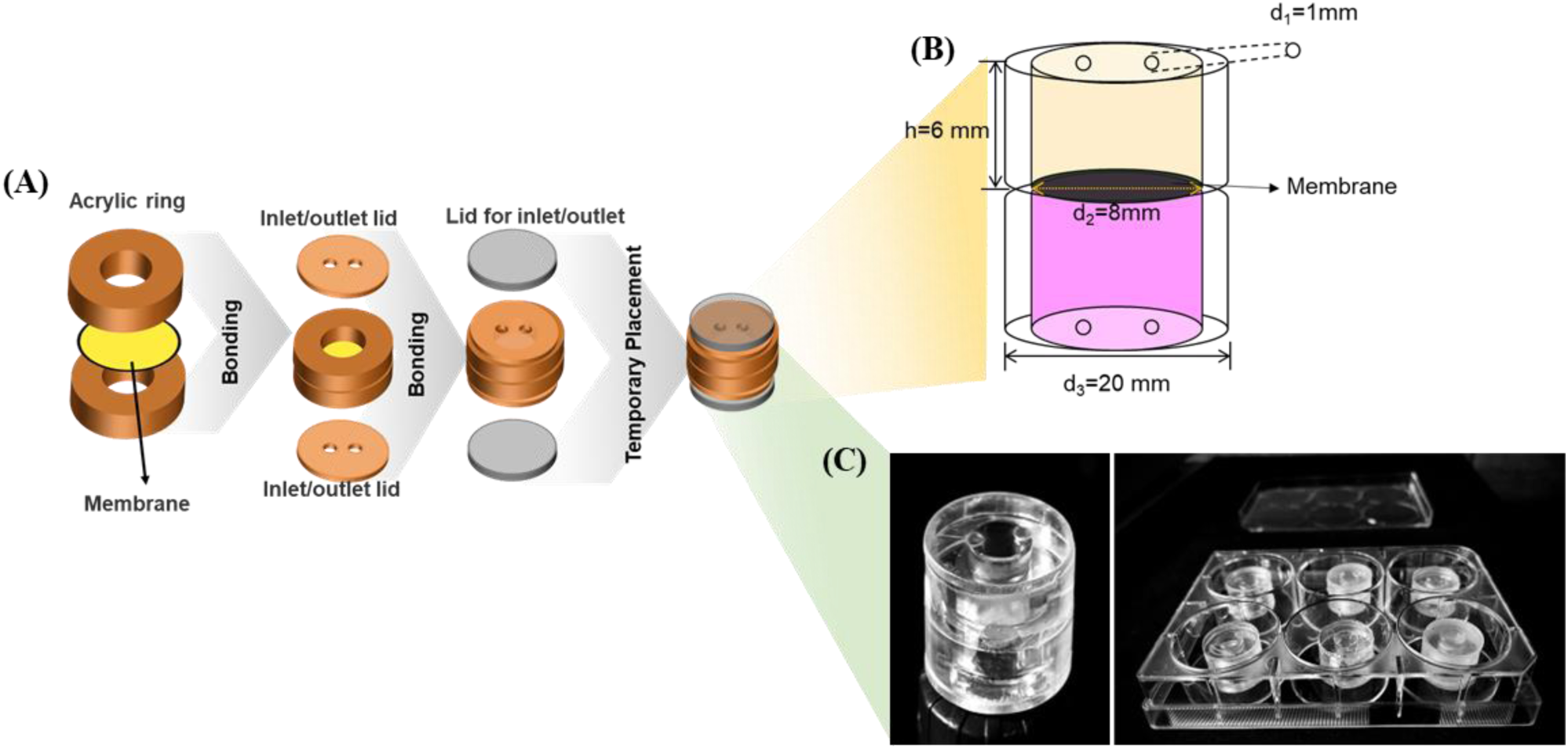
Device fabrication. (A) Schematic outlining the construction of a acrylic -based micro physiological device: PDMS stamp layers with defined inlet and outlet were bonded and temporarily assembled for experimental use. (B) Design of the cylindrical two-chamber insert showing key dimensions (C) Microphysiological placenta-on-a-chip prototypes device (left) and multiple devices positioned in a 6-well plate format for parallel experiments (right).

The device consists of two acrylic reservoirs bonded across a polycarbonate membrane (1 µm pore size), which serves as a structural analogue of the placental basal lamina (Figure 2A). The upper chamber represents the maternal compartment and supports trophoblast culture, whereas the lower chamber models the fetal compartment and accommodates endothelial cells. Independent access ports on both chambers enable cell seeding, medium exchange, and sampling without disturbing the opposing compartment. All device components were fabricated from laser-cut acrylic and assembled via PDMS-based stamp bonding, resulting in a robust, leak-proof construct suitable for repeated handling. Circular reservoirs with an inner diameter of 8 mm (d2) and an outer diameter of 20 mm (d3) were cut from 6 mm-thick acrylic sheets. Acrylic sheets of 1 mm thickness were cut to generate 20 mm diameter circular lids containing two 1 mm diameter (d1) ports, which were used as inlet and outlet points for cell seeding, medium exchange, and sample collection (Figure 2B). The 2 acrylic rings were separated by a polycarbonate membrane (Whatman Cyclopore, 1 µm pore size, 10 mm diameter) with an effective culture diameter of 8 mm, ensuring consistent transport area across replicates and enabling quantitative permeability measurements. These dimensions yielded a total volume of 300 µL. This volume is sufficient for downstream analysis, and even after the collection of 1 μL medium from the fetal chamber at different time points, the large volume ensures fluid stability. The overall dimensions were selected to allow the assembled devices to be housed within standard six-well plates, enabling parallel experiments and compatibility with conventional cell culture workflows. This configuration allows multiple devices to be operated simultaneously under identical conditions, supporting reproducibility and medium-throughput experimentation (Figure 2C).

### 3.2 Establishment of a trophoblast–endothelial bilayer interface

To evaluate whether the device supports the formation of a spatially organised maternal–fetal interface, BeWo trophoblasts and HUVEC endothelial cells were sequentially seeded on opposite sides of the semipermeable membrane. Prior to cell seeding, the membrane was coated with a thin layer of Matrigel to provide an extracellular matrix interface analogous to the placental basal lamina, facilitating uniform cell attachment on both sides of the membrane. Cell seeding densities were optimised to ensure uniform attachment and continuous layer formation on both surfaces of the membrane (Supplementary Table 1).

Fluorescent imaging of cell tracker–labelled cultures demonstrated the stable formation of two distinct cellular layers within the device (Figure 3A). BeWo cells populated the maternal-facing surface of the membrane, while HUVECs adhered uniformly to the fetal-facing side, with minimal cross-contamination between compartments. Higher-magnification views confirmed confluent coverage of the membrane by both cell types (Figure 3B).

**Figure 3.**
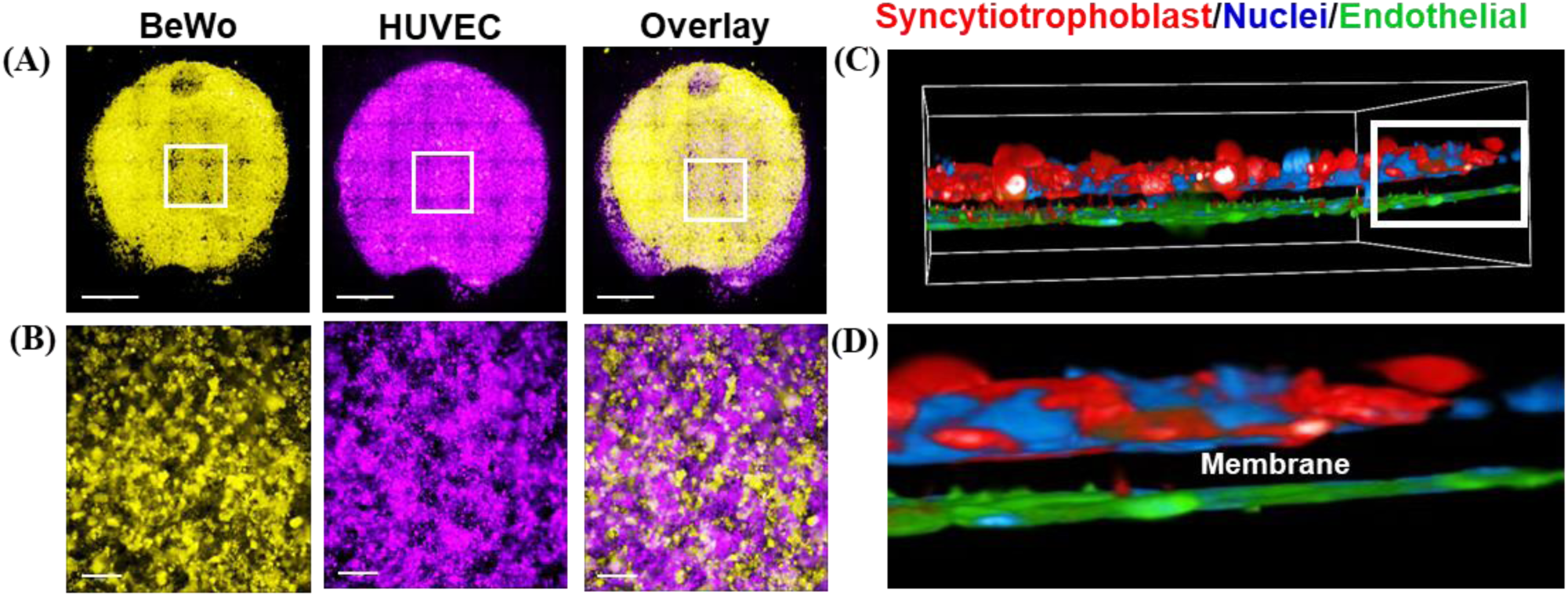
Co-culture of trophoblast and endothelial cells to reconstitute the placental barrier. (A) Fluorescent image showing confluent layers of BeWo and HUVECs on opposite sides of the interface. The cells were stained with Cell Tracker™ CMTPX (BeWo) and CellTracker™ CMFDA (HUVECs) respectively and pseudo coloured for representation. Scale bar 2 nun. (B) Zoomed image of the marked region horn (A), Scale bar 200 pm. (C) Three-dimensional visualization of the engineered placental barrier showing trophoblasts labeled with E-cadherin (CDH1) (red) and endothelial cells with VE-cadherin (CDH5) (green) with cell nuclei visualized by DAPI (blue). (D) Zoomed image of the marked region from (C). Marked region shows membrane between the 2 cell types is membrane.

To further assess the spatial organisation of the bilayer, three-dimensional imaging and orthogonal reconstructions were performed following immunostaining for cell-type–specific markers. BeWo trophoblasts exhibited E-cadherin (CDH1) expression on the maternal side of the membrane, whereas HUVECs expressed VE-cadherin (CDH5) on the opposing surface (Figure 3C). Z-stack reconstructions revealed two discrete cellular layers separated by the intervening membrane, confirming the successful reconstruction of a bilayer interface analogous to the placental barrier (Figure 3D).

Cell viability within the co-culture system was assessed using live–dead staining, which demonstrated high viability of both trophoblast and endothelial layers over the experimental time frame (Supplementary Figure 1). Together, these results establish that the device supports a robust co-culture of trophoblast and endothelial cells in a spatially defined configuration, providing a stable structural foundation for subsequent functional and transport studies.

### 3.3 Functional differentiation of trophoblasts into syncytial layer and endocrine activity

Following the establishment of a stable trophoblast–endothelial bilayer, we next evaluated whether the on-chip trophoblast layer could undergo functional differentiation into a syncytialised state, a defining feature of the human placental barrier. BeWo trophoblasts seeded on the maternal-facing surface of the membrane were treated with forskolin for 48h to induce syncytialisation before endothelial cell seeding (Figure 4A).

**Figure 4.**
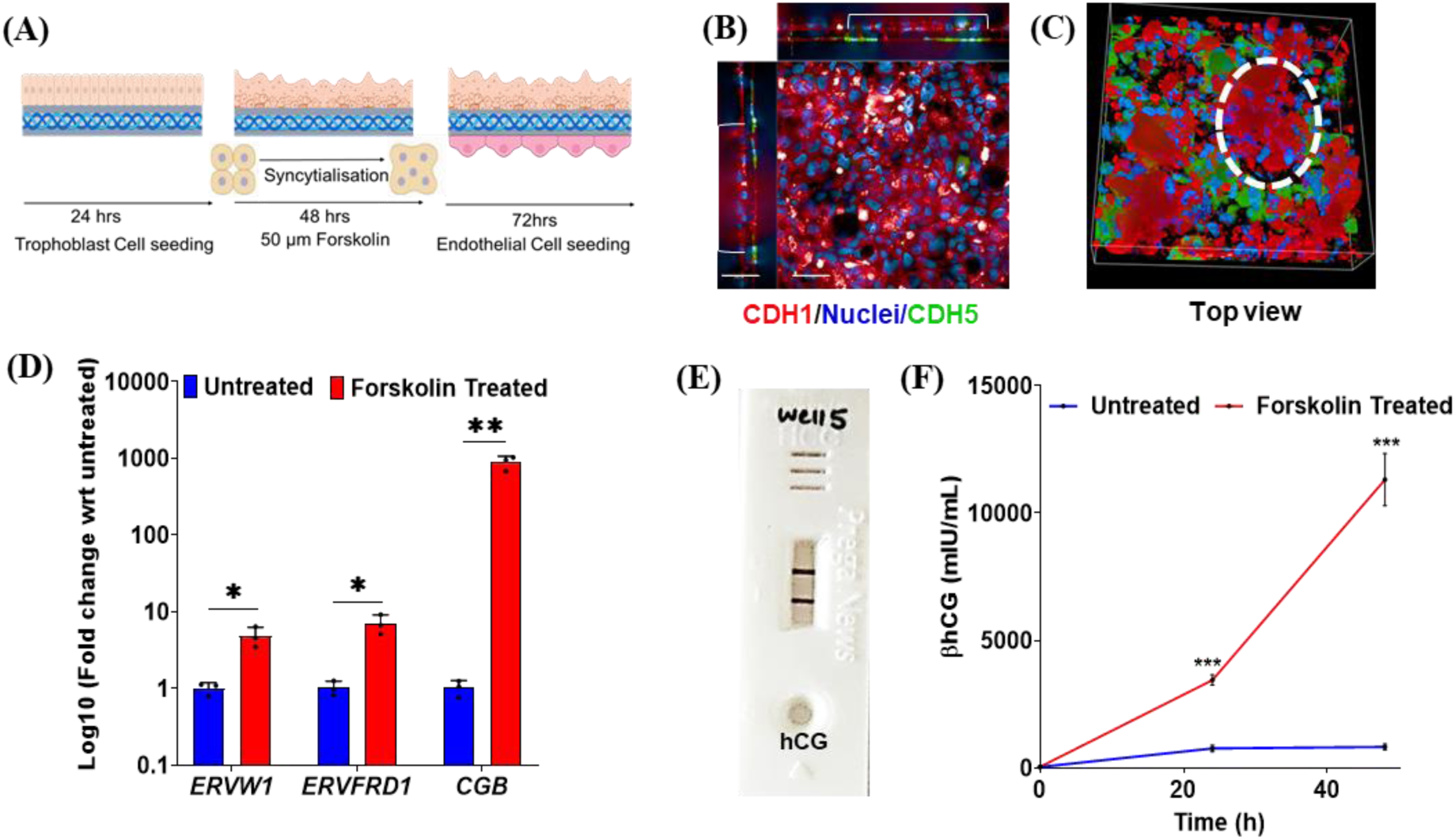
Svncytiotrophoblast differentiation and hormone production in the device. **(A)** Experimental illustr ation of the forskolin-induced syncytialisation in the device (B) 3D rendering side and top view of the trophoblast-fetal cell bilayer in the device, Scale bar 50 pm (C) Immunofluorescence images showing multinucleated syncytial structures in BeWo cells (red, syncytiotrophoblast regions); marked region shows Synctia. **(D)** Forskolin-induced syncytiotrophoblast differentiation. Y axis Log_lo_ fold change in mRNA expression of syncytialization markers *ERVJV-1, ERVFRD-1,* and *CGB* (P-hCG subunit) in forskolin-treated versus untreated trophoblast cells (n =3) Forskolin treatment significantly upregulated all markers. (E) Qualitative assessment of honnone production (phCG secretion). 2 purple lines in forskolin treated cell supernatant confirming honnone production and functional syncytiotrophoblast differentiation using pregnancy kit. (F) Qualitative assessment of phCG secretion using immunoassay showing increase in Forskolin treated cells. Y axis represents phCG concentration in mlU/mL; X axis represents time. Red line with black dots represents forskolin treated cells and blue with black dot represents untreated cells. * indicates pvalue <0.05, ** indicates pvalue <0.005, **** indicates pvalue<0.00005.

Immunofluorescence analysis revealed that following forskolin treatment, CDH1 exhibited a diffuse staining pattern, consistent with trophoblast fusion. Three-dimensional reconstruction further demonstrated the presence of contiguous, multinucleated regions spanning the membrane surface, indicating formation of a continuous syncytial layer within the device (Figure 4D-E). Importantly, this differentiated trophoblast layer remained stably integrated with the underlying endothelial compartment, preserving the bilayer architecture.

To determine trophoblast differentiation at the molecular level, expression of syncytialisation markers *ERVW-1*, *ERVFRD-1*, and CGB (β-hCG subunit) was assessed by quantitative PCR. The results revealed a significant increase in mRNA levels of *ERVW-1*, *ERVFRD-1*, and *CGB* in the Forskolin-treated group as compared to control (Figure 4D).

A key functional hallmark of syncytiotrophoblast differentiation is endocrine activity. Qualitative assessment using a rapid pregnancy detection assay confirmed the presence of β-human chorionic gonadotropin (β-hCG) in the culture medium collected from forskolin-treated devices (Figure 4B). Quantitative measurement demonstrated a significant increase in β-hCG secretion within 24h of forskolin treatment, with sustained levels observed over the experimental period (Figure 4C).

Together, these findings demonstrate that the placenta-on-chip platform supports trophoblast functional maturation into a hormone-secreting syncytial layer while maintaining structural integration with the endothelial compartment. This capability enables simultaneous assessment of placental barrier properties and endocrine function within a single, accessible in vitro system.

### 3.4 Evaluation of barrier integrity and selective permeability

A defining functional attribute of the placental barrier is its selective regulation of solute passage based on molecular size. To assess whether the engineered bilayer exhibits size-selective permeability, we examined the transport of a low molecular weight fluorescent tracer, fluorescein isothiocyanate (FITC ∼370 Da), and a high–molecular weight tracer, FITC–dextran (70 kDa), across the placenta-on-chip device (Figure 5A).

**Figure 5.**
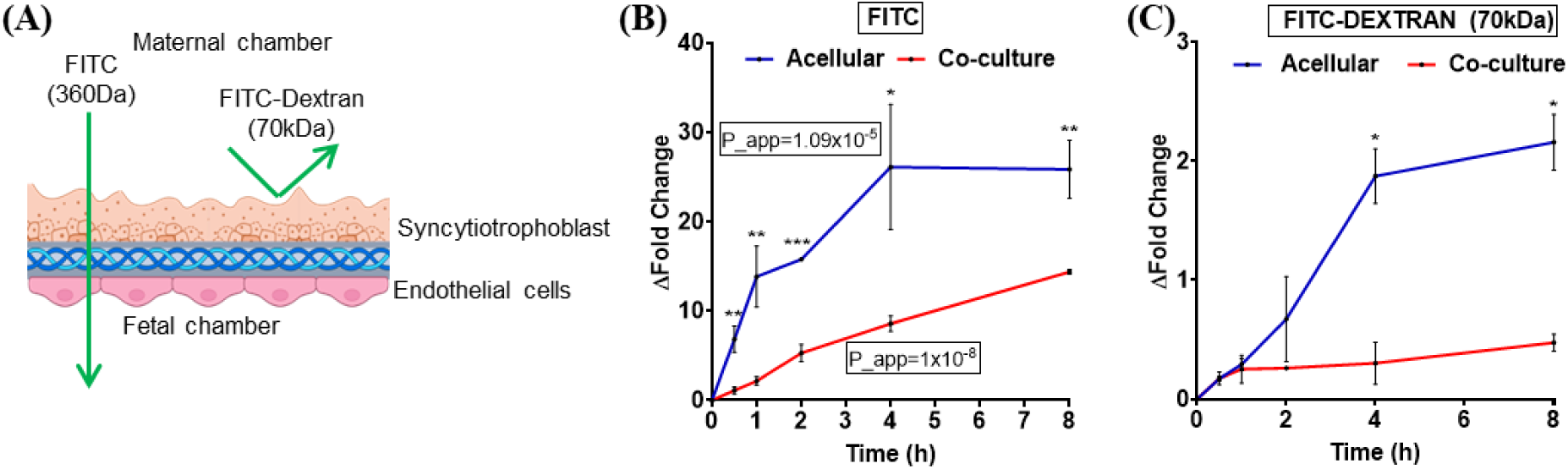
Barrier integrity of the placenta-on-chip device. **(A)** Illustration showing selective transport of FITC and FITC-Dextran across the placental barrier, (B) FITC permeability assay confirming reduced paracellular leakage across the barrier. Y-axis represents concentration of FITC in Fetal Chamber wrt T_o_and X-axis represents time. Each point represents value horn 3 devices with SD at particular time. Apparent permeability was calculated and represented in the graph (Acellulai- 1.09×10 ^5^ cm s ^1^ vs BeWo HUVEC= 1×10 ^8^ cm s’) (C) Quantitative analysis of barrier integrity showing limited transfer of FITC-Dextran (70kDa) Y-axis represents concentration of FITC- Dextran in Fetal Chamber wit T_o_and X-axis represents time. Each point represents value from 3 devices with SD at particular time. * indicates pvalue<0.05, ** indicates pvalue<0.005, *** indicates pvalue<0.0005.

In acellular devices containing the membrane alone, FITC rapidly diffused into the fetal chamber, with fluorescence levels increasing sharply during the initial hour and approaching equilibrium over the experimental time course (Figure 5B). In contrast, devices containing the trophoblast–endothelial bilayer exhibited a pronounced reduction in FITC transport. Although a gradual increase in fetal fluorescence was observed over time, the magnitude of FITC accumulation was substantially lower than that measured in acellular controls. Consistent with these observations, apparent permeability coefficients (P_app) calculated for FITC were reduced by a factor of 100 compared to the acellular condition. (Accelular= 1.09×10-5 cm s^-1^ vs BeWo HUVEC= 1×10-8 cm s^-1^) (Ratio cellular/acellular = 0.013) in the presence of the cellular bilayer, indicating restricted paracellular diffusion within the cellular barrier. (Figure 5B).

The barrier’s permeability to larger macromolecules was assessed using FITC–dextran. In membrane-only controls, FITC–dextran exhibited time-dependent accumulation in the fetal compartment, consistent with passive diffusion through the porous membrane. However, in devices containing the cellular bilayer, FITC–dextran transport was effectively suppressed, with fluorescence levels in the fetal chamber remaining near baseline throughout the experiment. No progressive increase in signal was observed at later time points, indicating sustained barrier integrity over time and absence of delayed leakage (Figure 5C)

Together, these results demonstrate that the placenta-on-chip model supports stable, size-selective permeability, permitting limited passage of small solutes while effectively restricting macromolecular transport. This selective barrier behaviour provides a functional foundation for subsequent evaluation of physiologically relevant metabolite transport across the engineered maternal–fetal interface.

### 3.5 Glucose transport across the placenta-on-chip under different culture configurations

Following validation of size-selective barrier behaviour, we next evaluated whether the placenta-on-chip model supports regulated transport of a physiologically relevant nutrient. Glucose transport was examined across devices configured as acellular controls, trophoblast monocultures, and trophoblast–endothelial co-cultures, allowing systematic assessment of the contribution of each cellular component to barrier function (Figure 6A–B). A handheld glucometer was used for glucose quantification due to its ability to measure glucose from microlitre-scale sample volumes, enabling repeated sampling from the same device for kinetic analysis. To ensure reliability of glucometer-based measurements, we performed a validation assay using a standard enzymatic hexokinase assay in a subset of samples, and observed a strong agreement across the tested range (R^2^=0.96), thereby validating our approach.

**Figure 6.**
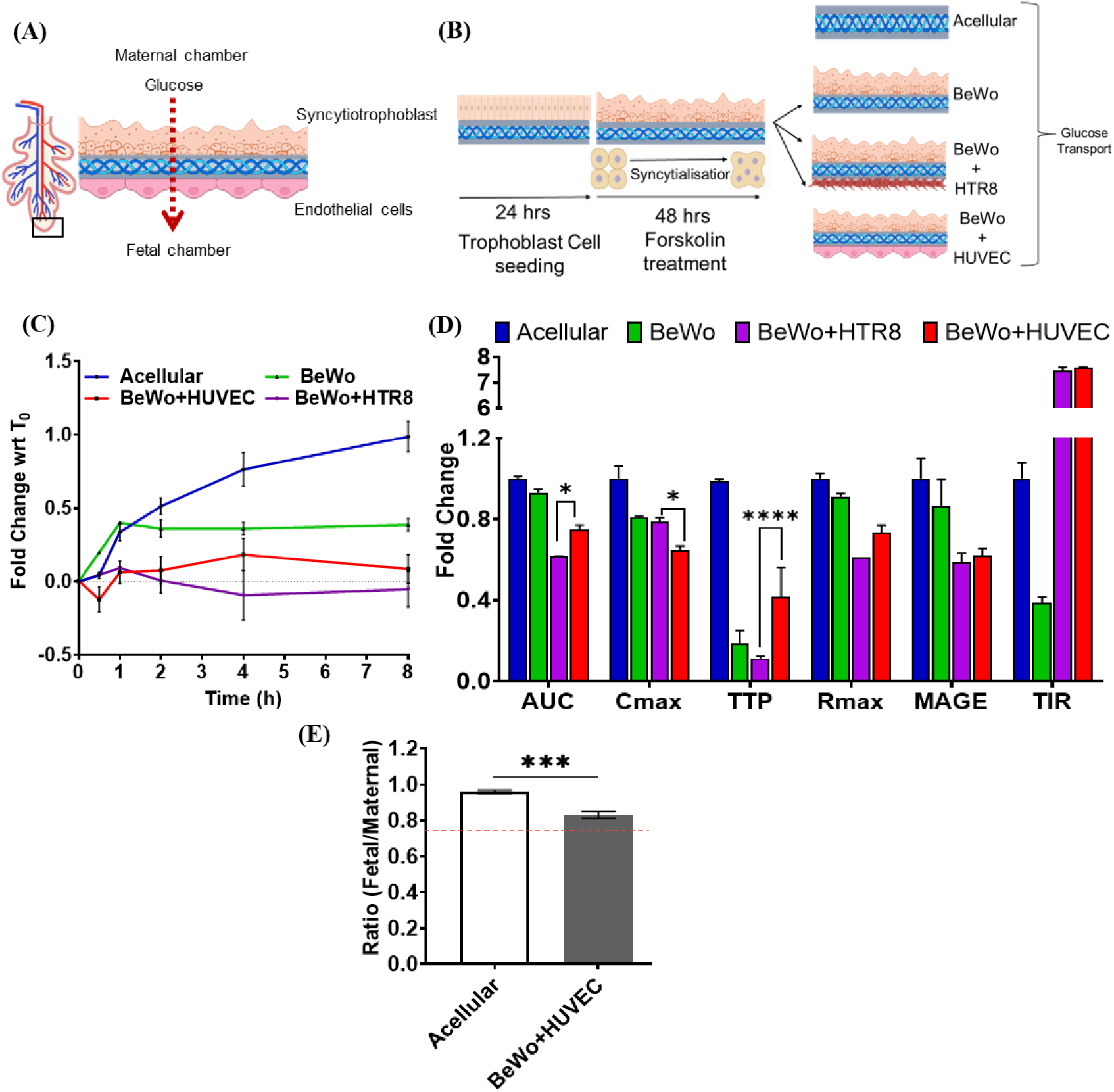
Evaluation of glucose transport across placenta-on-chip models under different culture configurations. (A) Schematic representation of matemal-to-fetal glucose transfer across the human placental barrier. (B) Experimental setup illustrating forskolin-induced syncytialisation of BeWo cells followed by culture in monoculture or co-culture with HUVEC or HTRS cells; acellular condition served as controls. (C) Time-dependent glucose transport in acellular, BeWo monoculture, BeWo+HTRS and BeWo+HUVEC conditions expressed as fold change relative to T_o_. Y-axis represents concentration of glucose in fetal chamber wrt T_o_ and X-axis represents time points. Each dot represents mean value from 3 devices with SD. (D) Pharmacokinetic parameters under acellular, BeWo monoculture, BeWo+HTR8 and BeWo+HUVEC conditions expressed as fold change relative to accelular . Bars represent Area Under the curve (AUC), Maximum Concentration (C,^, Time to Peak (TTP), Peak Rate (R_mfl_j, Mean Amplitude of Glycaemic Excursions (MAGE), and Tune in Range (TIR) for acellular, BeWo, BeWo+HTR8, and BeWo+HUVEC groups. (E) Comparison of fetal/matemal glucose ratios shows significantly reduced glucose transfer in the BeWo+HUVEC co-culture compared to the acellular. Dotted red line shows *in vivo* fetal/matemal ratio (0.75) (34). * indicates pvalue <0.05, ** indicates pvalue<0.005, *** indicates pvalue<0.0005, **** indicates pvalue<0.00005.

In acellular devices, glucose rapidly equilibrated between the maternal and fetal chambers, with fetal-to-maternal glucose ratios approaching unity within the experimental time frame, consistent with unrestricted diffusion across the membrane (Figure 6C). In contrast, devices containing the differentiated trophoblast layer exhibited a marked attenuation of glucose transfer. Although glucose levels in the fetal chamber increased over time, equilibrium was not reached, indicating the establishment of a functional barrier that modulates nutrient flux.

While differentiated trophoblast monocultures were sufficient to attenuate glucose diffusion, the placental barrier in vivo comprises both trophoblast and endothelial layers. We therefore next evaluated whether the incorporation of an endothelial compartment further modulates glucose transport dynamics. Compared with trophoblast monocultures, trophoblast–endothelial co-cultures displayed significantly reduced glucose accumulation in the fetal compartment and a sustained deviation from equilibrium over time (Figure 6C).

To further examine the specificity of endothelial contributions, glucose transport was compared across co-cultures comprising BeWo trophoblasts paired with either HUVECs or HTR8-SVneo cells. Whereas BeWo–HUVEC co-cultures exhibited gradual and regulated glucose transfer, BeWo–HTR8-SVneo co-cultures showed pronounced suppression of glucose transport, with fetal glucose levels remaining near baseline throughout the experiment (Figure 6C).

We next quantified glucose transport dynamics using integrated and peak-based parameters, including Area Under the Curve (AUC), maximum concentration (C_max_), time to peak (TTP), maximum transport rate (R_max_), and variability metrics such as mean amplitude of glycemic excursions (MAGE) and time in range (TIR), across acellular, BeWo monoculture, and co-culture configurations incorporating distinct endothelial cell types (Figure 6D). Relative to the acellular condition, cellular configurations exhibited clear alterations in transport behaviour across multiple parameters, consistent with barrier-mediated modulation of glucose flux (Supplementary Table 3). BeWo monoculture resulted in attenuation of glucose transfer, although integrated exposure (AUC) and peak transport rate (R_max_) were not markedly different from acellular conditions, indicating that trophoblasts alone impose partial, but not complete, restriction of transport. Importantly, the incorporation of an additional cell type produced a markedly different transport phenotype that depended on cell type (HTR8-SVneo vs HUVEC). Both co-culture models exhibited significantly reduced total glucose exposure (AUC), peak concentration (C_max_), and transport rate (R_max_) compared to acellular conditions, indicating establishment of a functional barrier. However, clear differences emerged between endothelial configurations. BeWo– HTR8-SVneo co-cultures demonstrated the greatest reduction in overall glucose exposure (AUC) and the lowest transport rates (R_max_), indicating a strong restriction of cumulative transport. In contrast, BeWo–HUVEC co-cultures exhibited the lowest peak glucose levels (C_max_) and the highest time in range (TIR), indicating enhanced stabilization of glucose dynamics despite a slightly higher cumulative exposure (AUC) compared to HTR8-based systems.

Variability analysis further supported these differences, with both co-culture systems showing reduced MAGE relative to acellular and BeWo-only conditions, but with distinct dynamic profiles (Figure 6D). BeWo– HTR8-SVneo configurations were characterised by stronger suppression of transport magnitude, whereas BeWo–HUVEC systems exhibited more controlled and stabilised glucose transfer over time. This highlights the importance of endothelial composition in shaping both the magnitude and temporal regulation of nutrient transfer across the maternal–fetal interface.

In vivo, maternal and fetal blood maintain an approximate ratio of 0.75, facilitating efficient glucose transfer [41] . In our device, fetal-to-maternal glucose ratios remained below unity and closely matched reported human placental values (0.82), indicating that incorporation of the endothelial compartment improves barrier-mediated regulation of glucose transport (Figure 6F).

Collectively, these findings demonstrate that the placenta-on-chip platform resolves configuration-dependent differences in glucose transport, with endothelial inclusion playing a critical role in regulating nutrient flux across the engineered maternal–fetal interface. This configurability enables systematic interrogation of how distinct cellular architectures influence placental transport behaviour within a controlled in vitro setting.

### 3.6 Directional urea transport across the placenta-on-chip

To examine whether the engineered placental barrier supports directional transport of small metabolic waste molecules, we evaluated urea transfer across the trophoblast–endothelial co-culture. Urea was introduced into the fetal chamber, and concentrations in both fetal and maternal compartments were quantified after 8 h of incubation (Figure 7A).

**Figure 7:**
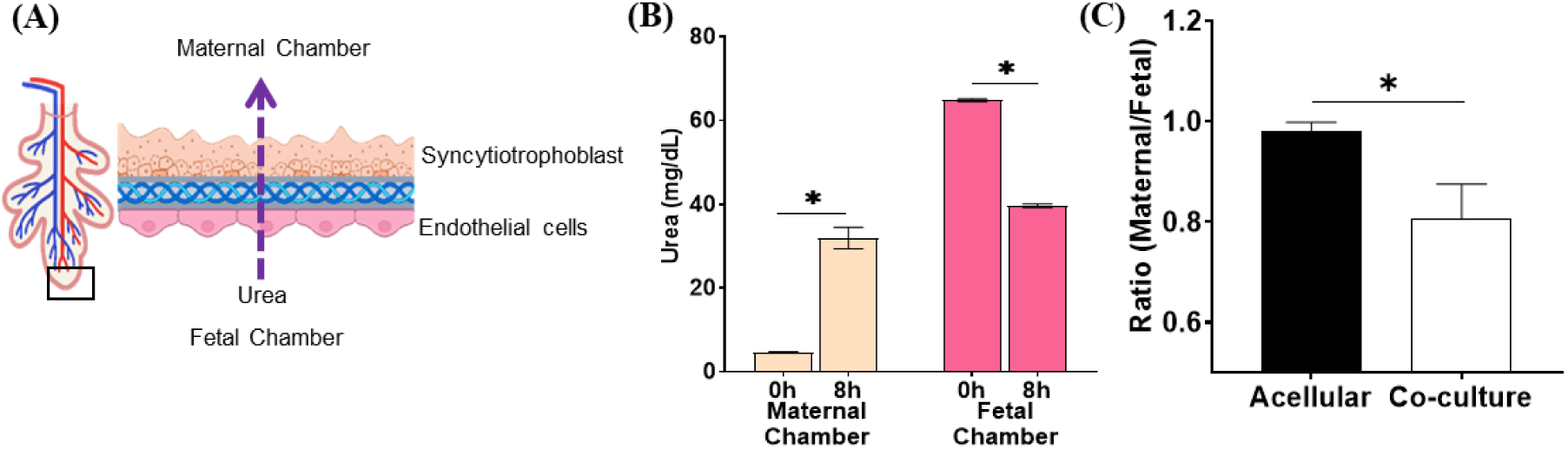
Urea Transport across the Placental Barrier. (A) Urea transfer kinetics demonstrating selective and physiologically relevant diffusion across the barrier. (B). Concentration of Urea in maternal and Fetal Chamber. Y-axis represents concentration of Urea (mg/dL) and X-axis represents time points from both maternal and fetal chamber. Each bar represents mean value from 5 devices with SD. (C) Comparison of matemal/fetal Urea ratio shows significantly reduction in urea transfer in the co-culture compared to the acellular. * indicates pvalue<0.05

At the 8 h endpoint, urea concentration in the maternal chamber was significantly increased compared with baseline, while a corresponding reduction was observed in the fetal chamber. This redistribution indicates effective transfer of urea across the bilayer from the fetal to the maternal compartment under static conditions (Figure 7B). Importantly, urea transfer occurred in the absence of overt barrier leakage, as the device maintained selective permeability characteristics established in prior assays.

To test if the cells contribute to any impedance to urea transport, we compared the ratio of urea in the maternal chamber to fetal chamber in the cellular and acellular conditions. The results revealed a small but significant reduction in the urea transported in the maternal chamber in the cellular conditions as compared to the acellular devices (Figure 7C)

These findings demonstrate that the placenta-on-chip model supports directional, small-molecule exchange consistent with physiological maternal–fetal waste transport, complementing the nutrient transport studies described above.

### 3.7 Altered glucose transport under hyperglycemic conditions

To evaluate whether the placenta-on-chip model can resolve changes in barrier behaviour under metabolic perturbation, glucose transport was examined in trophoblast–endothelial co-culture devices under normoglycemic (NG) and hyperglycemic (HG) conditions. Hyperglycemia was simulated by increasing glucose concentrations in the maternal compartment while maintaining constant fetal conditions (Figure 8A–B). Glucose transport profiles revealed distinct differences between acellular and co-culture systems under both conditions (Figure 8C). Under normoglycemia, acellular devices exhibited a gradual, monotonic increase in glucose levels over time, consistent with diffusion-driven transport. In contrast, co-culture devices showed attenuated glucose accumulation with early stabilisation, resulting in substantially lower glucose levels relative to acellular controls, indicating regulated transport across the cellular barrier. Under hyperglycemia, acellular systems displayed an increased magnitude of glucose transfer compared to normoglycemia, while retaining a similar monotonic profile, consistent with concentration-dependent diffusion. In contrast, co-culture systems exhibited a change not only in magnitude but also in the temporal pattern of glucose accumulation. Specifically, glucose levels increased rapidly during the initial phase, reached a peak at intermediate time points, and did not follow the sustained monotonic increase observed in acellular conditions (Figure 8C). These observations indicate that while hyperglycemia primarily amplifies diffusion-driven transport in acellular systems, it alters the temporal dynamics of glucose transport in the cellular model. This distinction between changes in transport magnitude and changes in transport dynamics forms the basis for the quantitative kinetic analysis described below.

**Figure 8:**
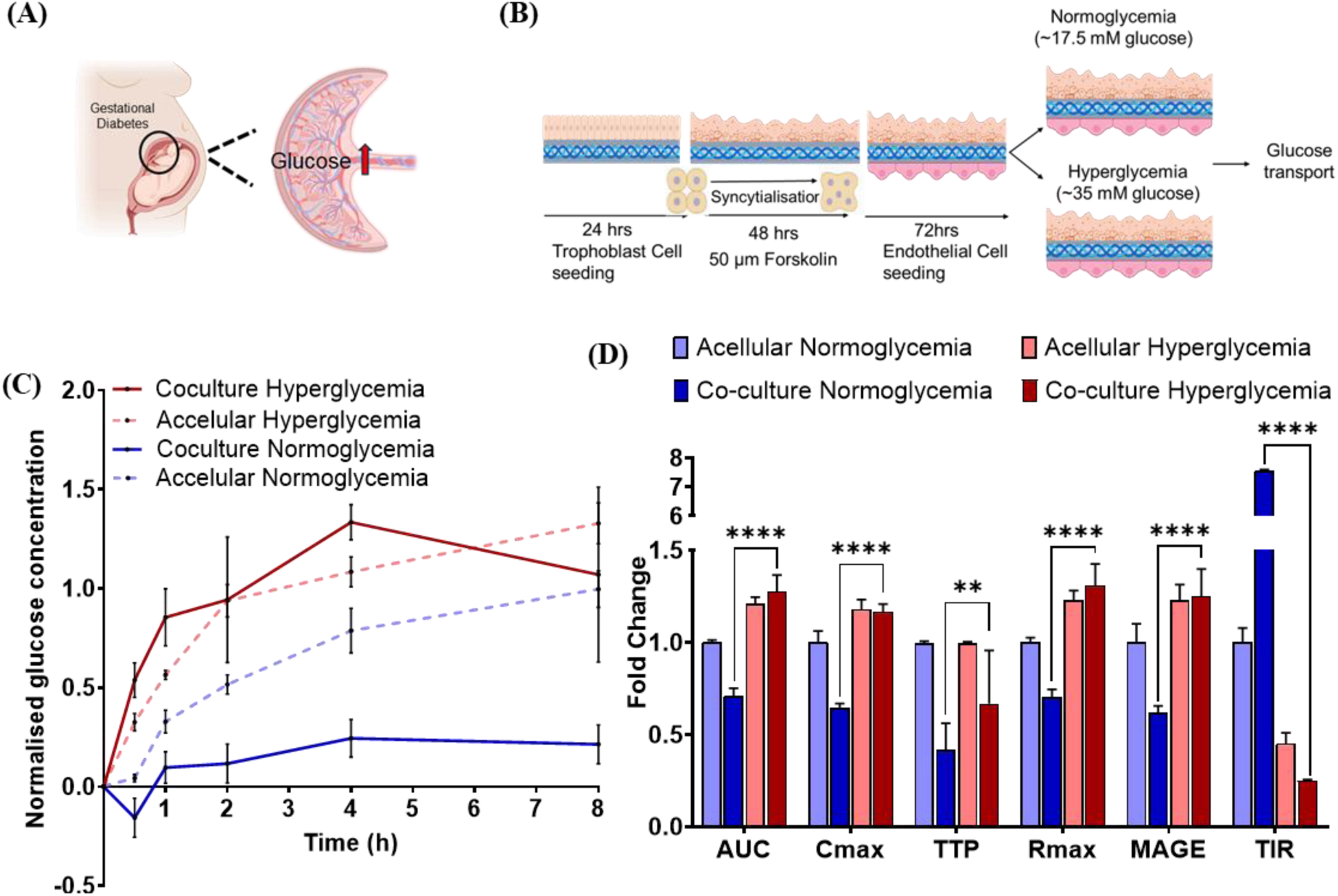
Kinetics of glucose transport during Hyperglycaemia. (A) Glucose transfer kinetics demonstrating selective and physiologically relevant diffusion across the barrier. (B) Experimental illustration of glucose transport across the placental barrier under normoglycemic (NG) and Hyperglycemic (HG) condition. (C) Glucose transfer measured in the fetal chamber. Y-axis represents concentration of glucose in fetal chamber normalized to T_o_ and X-axis represents time points. Each dot represents mean value from 3 devices with SD. Red solid and dashed line represents hyperglycemia co-culture and acellular conditions. Green solid and dashed line represents normoglycemia co-culture and acellular conditions. (D) Fold change in pharmacokinetic transport parameters. Bars show Area Under the curve (AUC), Maximum Concentration (C_max_), Time to Peak (TTP), Peak Rate (R^x), Mean Amplitude of Glycaemic Excursions (MAGE), and Time in Range (TIR) for acellular’ nonnoglycemia, acellular hyperglycemia, co-culture nonnoglycemia, and co-culture hyperglycemia .Each data point represents mean value from 3 devices with SD. * indicates pvalue<0.05; ** indicates pvalue<0.005, **** indicates pvalue<0.00005.

We also quantified glucose transport dynamics using integrated and peak-based parameters, along with variability metrics in normoglycemic (NG) and hyperglycemic (HG) conditions (Figure 8D and Supplementary Table 4). In acellular devices, glucose transport under hyperglycemia closely mirrored normoglycemic behaviour, with proportional increases in AUC, C_max_, and R_max_, while the temporal profile remained unchanged (TTP), consistent with diffusion-driven transport. In contrast, co-culture systems exhibited a distinct response to hyperglycemia. As compared to the normoglycemic condition, glucose transport increased significantly in magnitude (AUC, C_max_, R_max_). This was accompanied by increased variability (MAGE), reduced time in range (TIR), and a shift in time to peak (TTP), indicating altered regulation of glucose dynamics (Figure 8D).

Under hyperglycemic conditions, comparison between acellular and co-culture systems also revealed an interesting change in transport dynamics. AUC and R_max_ were modestly higher in co-culture systems compared to acellular controls, these differences were not statistically significant. In contrast, temporal and dynamic parameters differed significantly. Co-culture systems exhibited a markedly reduced time to peak (TTP) and a substantial decrease in time in range (TIR) relative to acellular conditions, indicating altered temporal regulation and reduced stabilization of glucose levels. These quantitative differences, together with the non-linear transport profiles observed in co-culture systems, support the conclusion that glucose transport under hyperglycemia is not governed solely by passive diffusion in the presence of cells.

Together, these findings demonstrate that the placenta-on-chip platform can resolve condition-dependent changes in both the magnitude and temporal regulation of nutrient transport, and can be used as a model to investigate metabolic perturbations such as hyperglycemia and to dissect their impact on placental barrier function.

## 4. Discussion

In this study, we present a static, two-chamber placenta-on-chip platform that recapitulates key structural and functional features of the human placental barrier in an experimentally accessible format. By co-culturing trophoblast and endothelial cells on opposing sides of a semipermeable membrane, the device reconstructs a maternal–fetal interface that supports trophoblast functional differentiation, selective solute permeability, and regulated nutrient and waste transport. Rather than seeking to replicate all aspects of placental physiology, the design prioritises functional barrier behaviour, open maternal compartmentalisation, and usability, providing a complementary alternative to more complex perfusion-based systems [42].

Several placenta-on-chip platforms have been developed that integrate trophoblast and endothelial compartments within microfluidic architectures, often incorporating perfusion to examine drug transfer and transport kinetics under controlled flow conditions [13–17,19–21,25,26,30–33,43–45]. A comparative overview of representative platforms and their design features is provided in Table 1. While these systems have generated important insights into placental transport behaviour, many emphasise microfluidic complexity and pharmacokinetic modelling, with comparatively less focus on trophoblast functional maturation and experimental scalability. A key physiological consideration is the haemochorial nature of human placentation, in which maternal blood directly bathes the syncytiotrophoblast surface within an open intervillous space [46,47]. Unlike vascularised organ systems, this interface experiences relatively low and discontinuous flow, and transport is predominantly diffusion-driven [1,22,44,47]. Estimates from human placental studies indicate that flow rates at the level of individual villous structures are on the order of ∼0.1–5 µL/min, corresponding to low shear conditions. In contrast, several microfluidic placenta-on-chip platforms operate at flow rates that are an order of magnitude higher (Table 1), thereby introducing shear environments that may not fully reflect the physiological maternal interface. Consequently, while perfusion-based systems are well suited for studying flow-dependent transport and pharmacokinetics, they may underrepresent diffusion-dominated exchange processes that are central to maternal–fetal transport of key metabolites such as glucose and oxygen. In this context, the present design provides a complementary framework that prioritises diffusion-driven transport within an open maternal compartment, while maintaining experimental simplicity and scalability. The static maternal configuration serves as a first-order approximation of the intervillous space for studying diffusion-driven transport, enabling physiologically relevant barrier formation without exposure to non-physiological shear. Within this architecture, we demonstrate stable formation of a trophoblast–endothelial bilayer and sustained barrier integrity across multiple functional assays. Importantly, the modular design can be extended to incorporate controlled fluid exchange or perfusion within the same geometry, allowing future integration of flow while preserving the core advantages of open maternal compartmentalisation, trophoblast differentiation, and plate-compatible operation.

A notable strength of the platform is its ability to support trophoblast functional maturation within the device. Like in 2D conditions[48], herein also we observed that forskolin-induced syncytialisation of BeWo cells resulted in the redistribution of E-cadherin and the formation of multinucleated syncytial regions, accompanied by robust β-hCG secretion in the on chip device. This ability to maintain endocrine activity alongside barrier function is a critical feature of the placental interface and is often underrepresented in the existing in vitro models, which focus primarily on permeability or transport kinetics [13–17,19–21,25,26,30–33,43–45]. By integrating trophoblast differentiation and hormone production into the same platform used for transport studies, the model enables simultaneous assessment of structural, functional, and endocrine aspects of placental biology.

Functional validation of the barrier was achieved through a combination of size-selective permeability assays and physiologically relevant metabolite transport studies. The trophoblast–endothelial bilayer effectively restricted macromolecular passage while permitting limited diffusion of small solutes, demonstrating stable and selective barrier behaviour. We further demonstrate that the glucose transport across the device was strongly dependent on cellular configuration, with endothelial cell inclusion significantly modulating both the magnitude and temporal dynamics of nutrient flux and yielding fetal-to-maternal ratios comparable to those reported in vivo [41]. In addition, the platform supported directional transfer of urea from the fetal to the maternal compartment, confirming its ability to model bidirectional exchange processes relevant to placental waste clearance.

Beyond baseline physiological function, the placenta-on-chip was responsive to metabolic perturbation. During pregnancy, maternal hyperglycemia is associated with increased glucose transfer across the placenta, contributing to fetal hyperglycemia and macrosomia [49–51]. In the present model, hyperglycemic conditions resulted in a marked increase in glucose transport across the barrier. While glucose transport under hyperglycemia increased in magnitude the temporal characteristics of transport were significantly altered relative to normoglycemic conditions. Specifically, co-culture systems showed changes in time to peak and a marked reduction in time in range, accompanied by increased variability in glucose levels. These findings indicate that hyperglycemia alters the temporal regulation of glucose transport in the cellular system rather than simply amplifying passive diffusion. Glucose transport in the placenta is primarily mediated by GLUT1 transporters expressed on trophoblast cells, including BeWo cells as established in prior studies [22,29,32,33]. While the present study focuses on functional transport outcomes, future investigations examining transporter expression and localisation under normoglycemic and hyperglycemic conditions will be important to elucidate the molecular mechanisms underlying these observations. Nevertheless, our observation positions the platform as a useful tool for studying how maternal metabolic states influence placental transport dynamics, without requiring complex flow control or advanced microfluidic infrastructure.

Design choices in biofabricated model systems inevitably reflect a balance between physiological fidelity, experimental robustness, and practical usability. In the present platform, both material selection and cell system choice were guided by the need for reproducible barrier formation and scalable functional assessment. While polydimethylsiloxane (PDMS) is widely used in microfluidic devices, its mechanical deformability, susceptibility to leakage during repeated handling, and non-specific adsorption of small molecules can complicate transport studies and limit reproducibility [52,53]. The use of rigid acrylic components enabled stable compartmentalisation, compatibility with standard multi-well plates, simplified operation, and supported parallel experimentation under controlled conditions. Similarly, although primary placental cells offer valuable biological insight, their limited availability, donor-to-donor variability, and inherent heterogeneity pose challenges for reproducibility and large-scale studies. In contrast, the use of well-characterised trophoblast and endothelial cell lines facilitates consistent differentiation, stable barrier formation, and comparative analysis across devices. Together, these design choices position the platform as a robust and industry-compatible system for functional placental barrier studies where reproducibility and scalability are prioritised. While the static configuration of the present platform reflects the diffusion-dominated maternal interface of human placentation, the absence of flow in the endothelial compartment may limit the ability to capture shear-dependent endothelial responses. Incorporation of controlled perfusion in the fetal channel within the same device geometry represents a potential future extension to further enhance its physiological relevance.

The modular and scalable nature of the placenta-on-chip platform enables a variety of future applications. The device can be adapted for evaluating transplacental drug transfer and barrier effects, supporting classification or reclassification of compounds based on placental permeability and barrier integrity, with potential relevance for pregnancy-safe drug development. The system also provides opportunities to identify modulators of nutrient transport, study placental responses to infectious agents, and investigate pathological conditions such as gestational diabetes and other pregnancy-associated disorders. The platform offers a controlled experimental framework for addressing fundamental questions in placental biology related to trophoblast–endothelial interactions and barrier regulation.

## 5. Conclusion

In summary, this work introduces a robust and accessible placenta-on-chip platform that complements existing perfused systems by emphasising open maternal compartmentalisation, functional trophoblast maturation, and experimentally tractable barrier assessment. By combining biological relevance with fabrication simplicity, the model provides a practical framework for studying placental barrier function, nutrient and waste transport, and metabolic perturbations, and may serve as a useful tool for both fundamental and applied investigations in maternal–fetal biology.

## Acknowledgements

We are thankful to Dr Bhakti Pathak (ICMR-NIRWoH) for sharing BeWo cells. This study was funded by ANRF-DST-IMPRINT II C grant (Anusandhan National Research Foundation, Department of Science and Technology; IMP/2019/000115). DM lab is supported by grants from Indian Council of Medical Research (ICMR) and Wadhwani Research Centre for Bioengineering (WRCB), IIT Bombay for funding and support. AB is the recipient of Council for Scientific and Industrial Research (CSIR)- Senior Research Fellowship (111-3629-1491/2K24/1). ChatGPT-5.2 was used to assist with language editing of the manuscript, under the authors’ supervision. The manuscript bears the ICMR-NIRWoH ID: OA/4/02-2026.

