## Supplementary Figure 1 for "A Functional Placenta-On-Chip Model For Maternal–Fetal Transport"

**Supplementary figure 1: Viability assessment of bi-layer**

**Description:** Viability assessment of endothelial cells and trophoblast cells in co-culture using Calcein/Hoechst staining

Supplementary figure 1: Viability assessment of endothelial cells and trophoblast cells in co-culture. A) Calcein/hoescht stained BeWo cells in the device B) Calcein/hoescht stained HUVEC cells in the device. Scale bar is of 400 µm at 10X. Green shows calcien stained cells. Hoechst shows nuclei. Scale bar is 1000 µm at 4X.

**Hoechst**

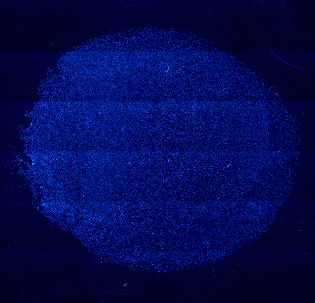

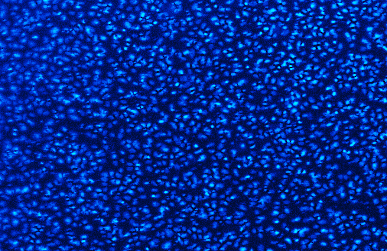

**Calcein**

**Scanned**

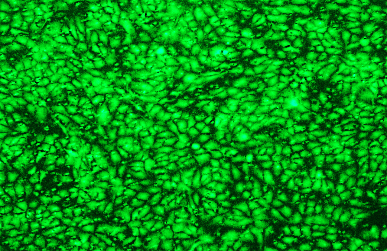

**10X**

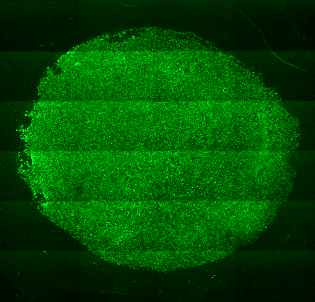

(B)

**HUVEC cells**

**Calcein**

**Hoechst**

**10X**

**Scanned**

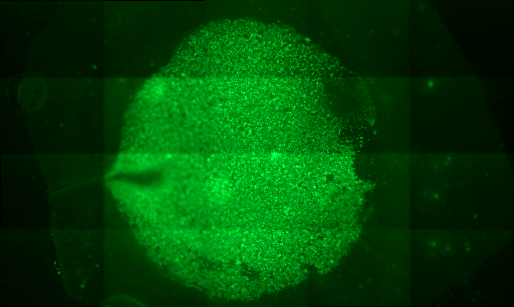

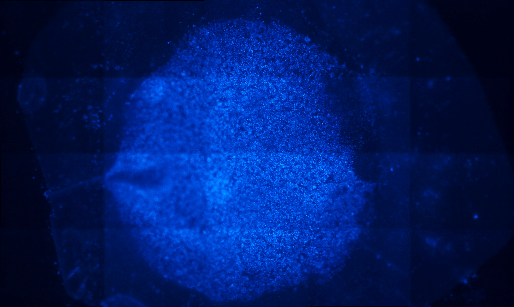

(A)

**BeWo cells**

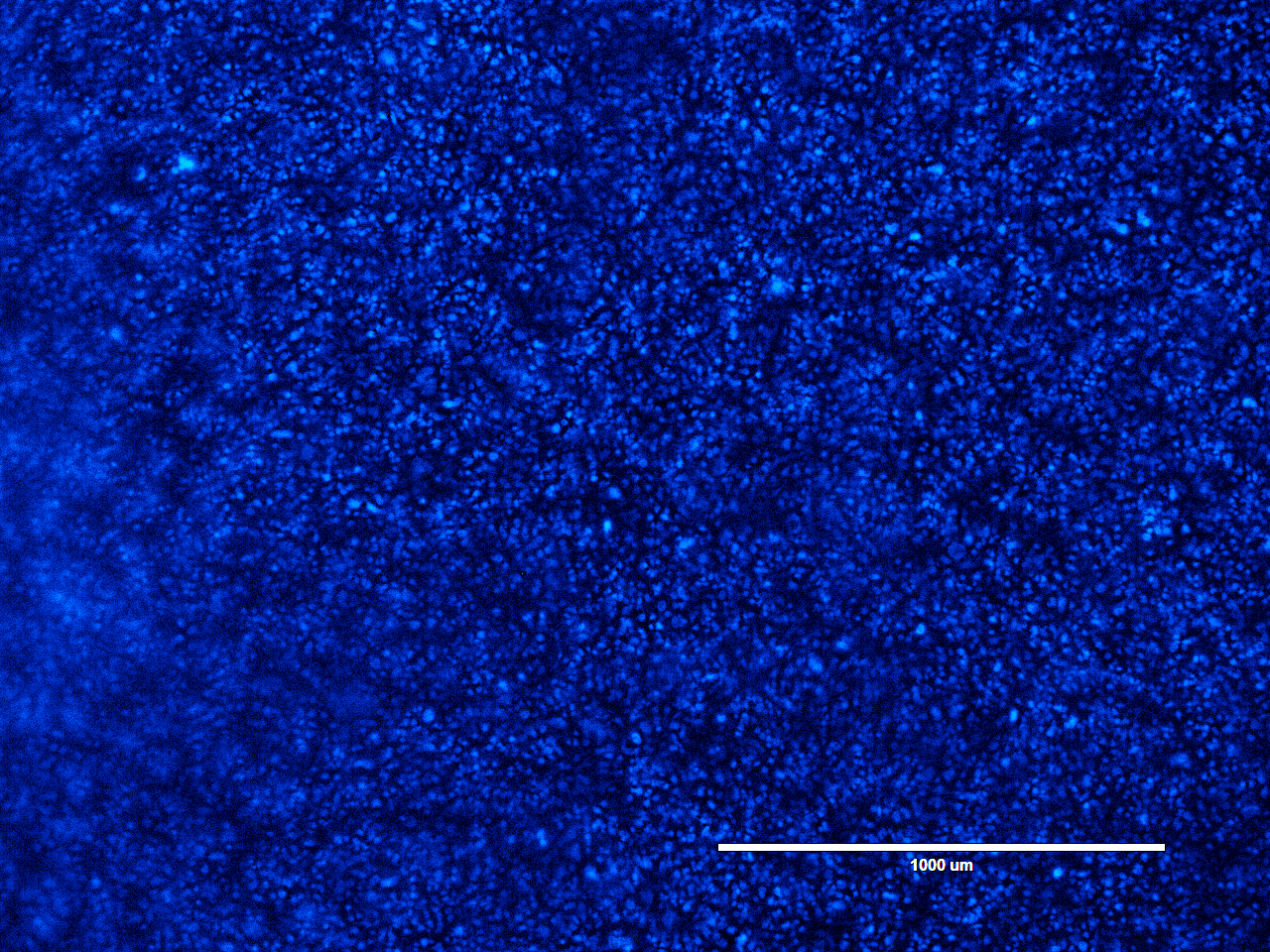

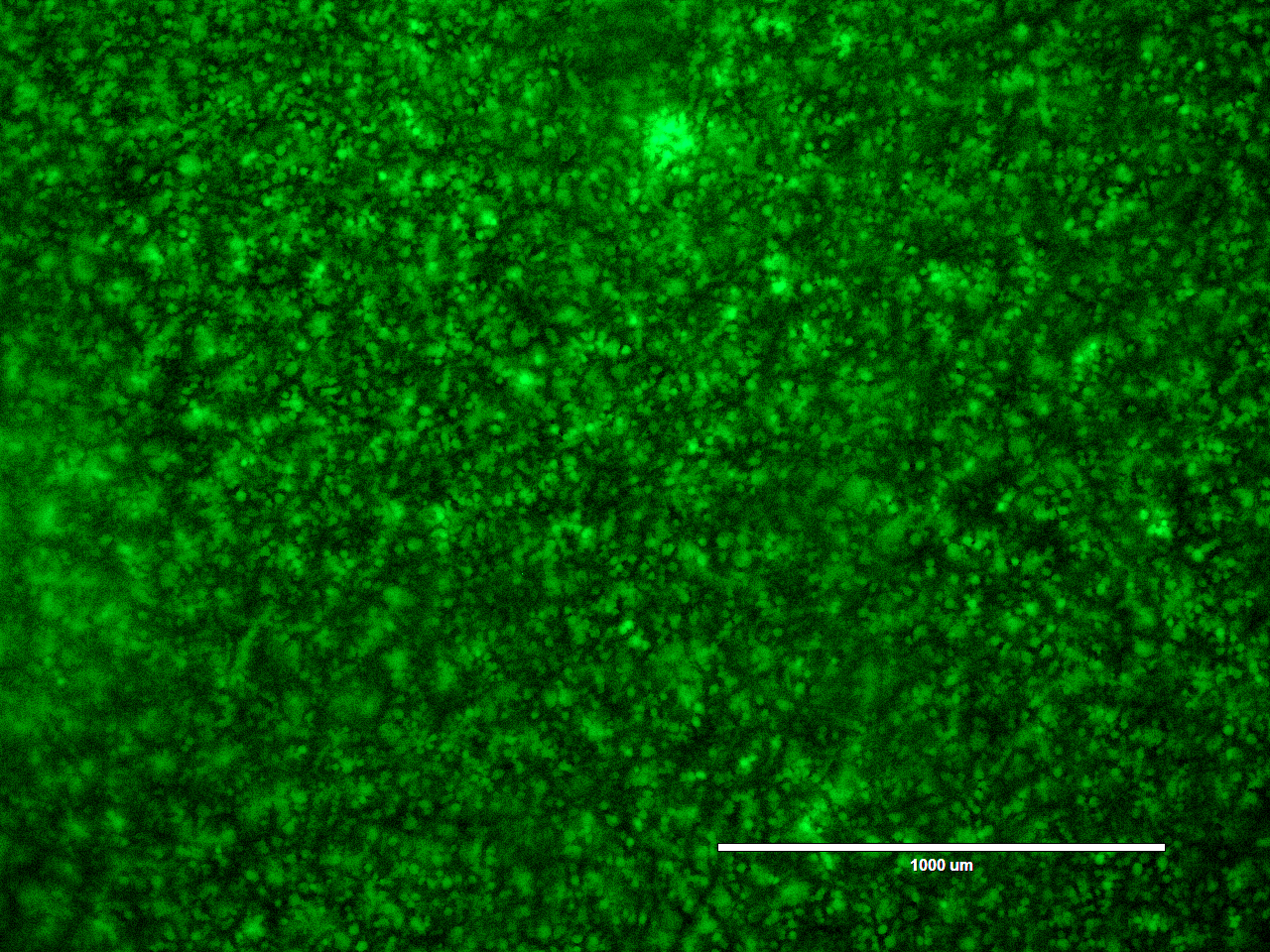
