## Supplementary Figure 2 for "A Functional Placenta-On-Chip Model For Maternal–Fetal Transport"

**Supplementary Figure 2: Standard curve for FITC and FIT- Dextran.**

**Description:** Standard curve for FITC and FITC-Dextran was made using Qubit 4. Fluorescence intensity was plotted as Log_10_ relative fluorescence units (RFU). FITC (Stock conc.- 264.5 µM) and FITC-Dextran (Stock conc.- 1.42 µM)) concentration dilutions 1:1,1:2,1:4,1:6, 1:8. Linear regression fit to the experimental data points was done.

Supplementary Figure 2: Standard curve for FITC and FIT- Dextran. (A) Standard curve for FITC was made using Qubit 4. Fluorescence intensity was plotted as Log_10_ relative fluorescence units (RFU) (Y-axis) against concentration dilutions 1:1,1:2,1:4,1:6, 1:8 (X-axis) from concentration of 264.5 µM. The red dotted line represents the linear regression fit to the experimental data points (R^2^=0.95). The FITC standard curve showed linear relationship between concentration dilution and Log_10_ RFU, indicating that fluorescence intensity decreased progressively with increasing dilution. (B) Standard curve for FITC-Dextran was made using Qubit 4. Fluorescence intensity was plotted as Log_10_ relative fluorescence units (RFU) against concentration dilutions 1:1,1:2,1:4,1:6, 1:8 (X-axis) from concentration of 1.42 µM. The red dotted line represents the linear regression fit to the experimental data points (R^2^=0.95). The FITC-Dextran standard curve showed a clear inverse, linear inverse relationship between concentration dilution and Log_10_ RFU, indicating that fluorescence intensity decreased progressively with increasing dilution. Error bar shows ±SD.

(A)

(B)

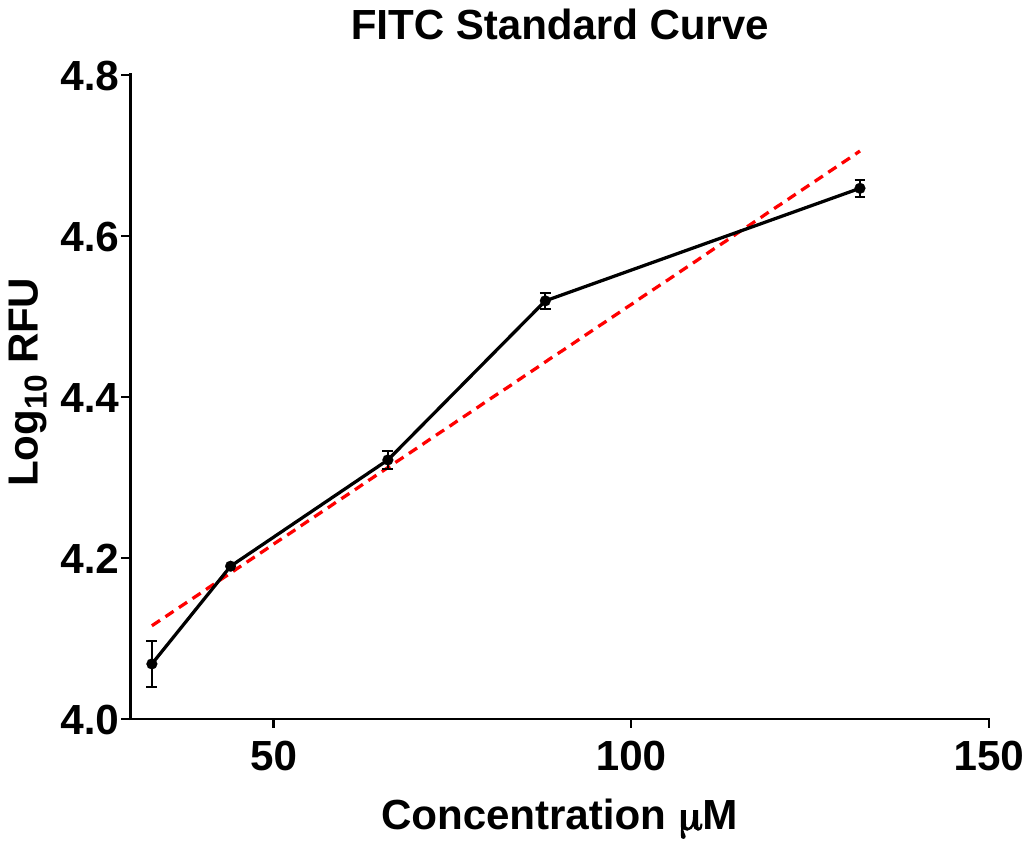

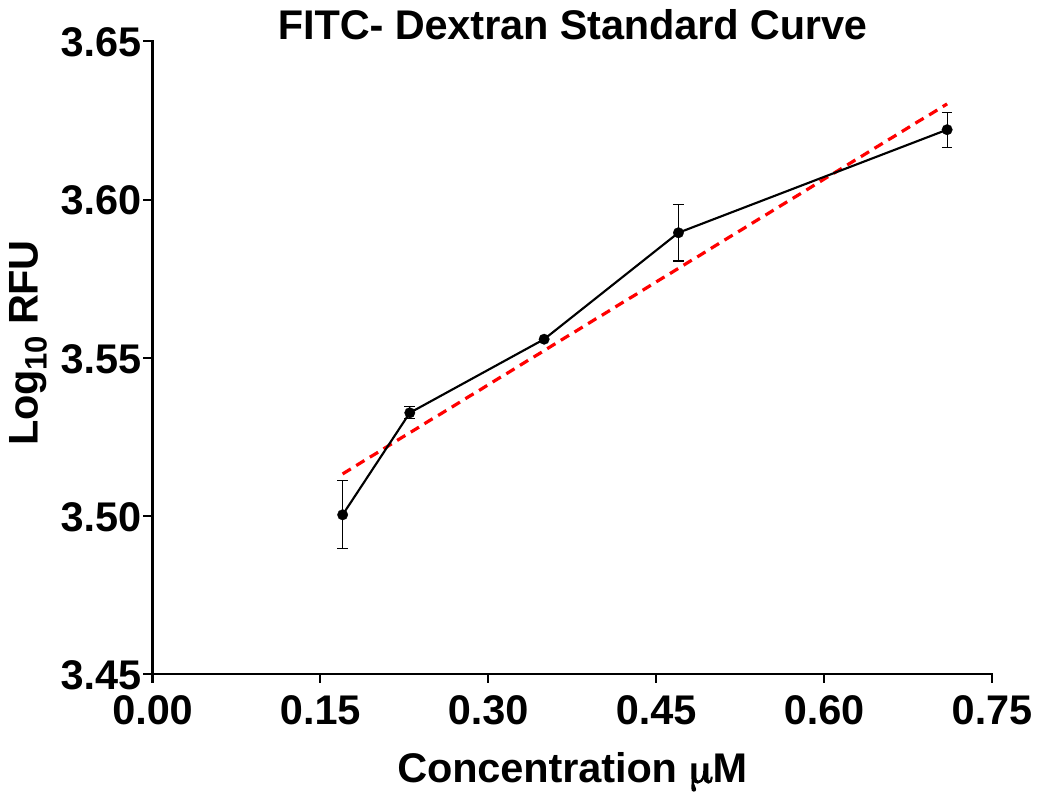
