## Supplementary Table 1 for "A Functional Placenta-On-Chip Model For Maternal–Fetal Transport"

**Supplementary Table 1: Extracellular Matrix (ECM) and cell seeding densities**

| *Cell type* | *ECM-Coating* | *Cell Seeding Density* | *Seeding conditions* |
| --- | --- | --- | --- |
| BeWo | Collagen | 0.75*10^5 | Overnight incubation |
|  | Collagen | 1*10^6 | Overnight incubation |
|  | Collagen | 1.5*10^6 | Overnight incubation |
| BeWo | Matrigel | 1*10^6 | Overnight incubation |
|  | **Matrigel** | **1.5*10^6** | **Overnight incubation** |
| HUVEC | Matrigel | 0.75*10^5 | Overnight incubation |
|  | **Matrigel** | **1*10^6** | **Overnight incubation** |
|  | Matrigel | 1.5*10^6 | Overnight incubation |

**Description:** Cell seeding densities for BeWo and HUVEC. Different cell number and extracellular matrix coating for the membrane were tested. Text in bold shows final used cell number and extracellular matrix coating used for final experiments
