## Supplementary Table 2 for "A Functional Placenta-On-Chip Model For Maternal–Fetal Transport"

**Supplementary Table 2: Primer sequences real-time PCR analysis**

**Description:** Primer sequences used for quantitative real-time PCR analysis of trophoblast differentiation marker genes, *ERVW1, ERVFRD1,* and *CGB.* Forward and reverse primer sequences together with annealing temperatures are listed.

| Gene | Primer Sequence | Annealing Temperature |
| --- | --- | --- |
| *18S* | Forward -5’ GGAGAGGGAGCCTGAGAAAC 3’  Reverse- 5’ CCTCCAATGGATCCTCGTTA 3’ | 60 |
| *ERVW1* | Forward -5’ATTGGCGGTATCACAACCT3’  Reverse- 5’ CGTGAGAATGAGAACCAG 3’ | 60 |
| *ERVFRD1* | Forward -5’CCACCAACATCCTTTCAA3’  Reverse- 5’ CCAGTGTTTCGAAGCTCCT 3’ | 54 |
| *CGB* | Forward -5’ CAGCATCCTATCACCTCCTGGT3’  Reverse- 5’ CTGGAACATCTCCATCCTTGGT 3’ | 62 |
